# Dynamics of contrast decrement and increment responses in human visual cortex

**DOI:** 10.1101/425173

**Authors:** Anthony M. Norcia, Alexandra Yakovleva, Bethany Hung, Jeffrey L. Goldberg

**Affiliations:** Department of Psychology, Stanford University; Department of Ophthalmology, Stanford University; Department of Psychology, Brown University

**Keywords:** Luminance contrast, ON, OFF pathway, Visual Evoked Potentials

## Abstract

The goal of the present experiments was to determine whether electrophysiological response properties of the ON and OFF visual pathways observed in animal experimental models can be observed in human. Visual Evoked Potentials (VEPs) were recorded in response to equivalent magnitude contrast increments and decrements presented using sawtooth temporal waveforms at a temporal frequency of 2.73 Hz. VEP response waveforms and response spectra for incremental and decremental stimuli were analyzed as a function of stimulus size and visual field location in 68 healthy adult participants. VEP response were larger in amplitude and shorter in latency for contrast decrements than for contrast increments. The spatial tuning was narrower for contrast decrements than for contrast increments and responses were larger for displays that were scaled for cortical magnification. VEPs recorded at the scalp differ between contrast decrements and increments of equal Weber contrast in a fashion that parallels results from the early visual system of cats and monkeys. Because our assay allows differential detection of ON and OFF pathway activity in human, the approach may be useful in future work on disease detection and treatment monitoring.

## Introduction

In the vertebrate retina, two parallel pathways diverge from the first synapse between the photoreceptors and bipolar cells: one signaling luminance increments (ON), and the other signaling luminance decrements (OFF) (Dowling and Werblin, 1971). These dual pathways flow from bipolar to retinal ganglion cells (RGCs) whose afferents stratify in different retinal sublaminae (Famiglietti and Kolb, 1976; Wassle, 2004), and they remain segregated in the lateral geniculate nucleus (LeVay and McConnell, 1982; Stryker and Zahs, 1983). Visual cortex also retains a degree of pathway segregation (McConnell and LeVay, 1984; Zahs and Stryker, 1988; Jin et al., 2011a; Smith et al., 2015; Wang et al., 2015; Kremkow et al., 2016).

The spatial properties of the two pathways have been extensively documented. The receptive fields of OFF RGCs are smaller than those of their ON counterparts (Dacey and Petersen, 1992; Devries and Baylor, 1997; Chichilnisky and Kalmar, 2002; Nichols et al., 2013), as are their dendritic arbors (Dacey and Petersen, 1992). OFF RGCs are also more numerous (Ratliff et al., 2010) and the cortical tiling of OFF inputs is more focal than ON tiling (Kremkow et al., 2016; Lee et al., 2016). OFF responses dominate ON responses in cortex (Jin et al., 2008; Yeh et al., 2009; Xing et al., 2010) and the contrast response function also differs for increments and decrements (Zaghloul et al., 2003). Together, these factors may explain the reported higher spatial resolution for darks than lights (Kremkow et al., 2014). Prior work with natural image statistics suggests the adaptive value of this asymmetry. Negative (dark) contrasts are more prevalent than positive (light) contrasts in natural scenes (Ratliff *et al*., 2010; Cooper & Norcia, 2015)(Geisler, 2008).

Differences in the dynamics of the two pathways have been less studied. An early in vitro study in primate RGCs found faster responses in ON *vs* OFF cells (Chichilnisky and Kalmar, 2002). This work was cited as a possible basis for an apparent motion illusion where it was estimated that brights were processed ~ 3 msec faster than darks (Del Viva et al., 2006). More recent in vivo work in cat has found that OFF cells in the LGN (Jin et al., 2011b) and in visual cortex (Komban et al., 2014) respond more quickly than ON-dominated cells. This latter physiological ON/OFF asymmetry is consistent with other human psychophysical work that has found that darks are processed faster than lights (Komban, Alonso, & Zaidi, 2011).

Visual evoked potentials (VEPs) provide a possible means of measuring the response properties of the two pathways non-invasively. Several VEP studies that have compared responses to contrast increments and decrements (Zemon et al., 1988; Mutlukan et al., 1992; Zemon et al., 1995; Roveri et al., 1997; Kremkow et al., 2014) on the assumption that luminance decrements are preferentially processed by the OFF pathway and vice versa (Kremers et al., 1993). Here we build on this prior research by presenting sawtooth increments and decrements using a spatially optimized stimulation array and large study samples. Our measurements indicate that VEPs to decrements are typically larger and faster than those to corresponding increments.

## Materials and Methods

### Participants

A total of 68 adults between the ages of 18 and 50 participated. All participants had visual acuity of 20/25 or better in each eye on the Bailey-Lovie constant LogMAR chart and a stereoacuity of 50 arc sec or better on the RandDot test.

### Visual Stimuli

We report the results of three experiments. The first experiment was designed to optimize the spatial stimulus parameters, the second was designed to obtain full-field data from a large group of participants and the third to compare response properties between the upper and lower visual fields.

We used periodic decremental and incremental sawtooth stimulation to elicit SSVEPs biased to OFF *vs* ON pathways, respectively (Kremers et al., 1993). Decremental stimuli were defined as those in which the fast phase of the sawtooth decreases in luminance and the slow ramp phase increases (Fig. 1A, red curve), with incremental stimuli being the opposite (Fig. 1A, black curve). The spatial structure of the stimulus was based on the Westheimer sensitization paradigm (Westheimer, 1965, 1967). In our version of the paradigm, a small probe presented on a larger background pedestal, was modulated according to either a decremental or incremental sawtooth profile just described (Fig. 1B). The probes were presented at a fixed modulation depth of 20% Weber contrast for both increments and decrements. Contrast was calculated as the probe luminance minus the pedestal luminance over the pedestal luminance. Increments and decrements had opposite signs but equal values in under this definition. Stimuli were presented on a SONY PVM-2541 monitor (1920×1080 pixels) viewed monocularly at a distance of 70 *cm*. The stimuli comprised multi-element arrays where each element comprised a probe-on-pedestal element (see Fig. 1C for an example). Pedestal luminance was 50 cd/m2, background luminance was 11 cd/m2 and probe contrast relative to the pedestal was +/−20% based on the Weber definition Lmax-Lmin/Lmin. Display pipeline delays and EEG pipeline delays were measured with a photocell and have been corrected.

**Fig. 1.**
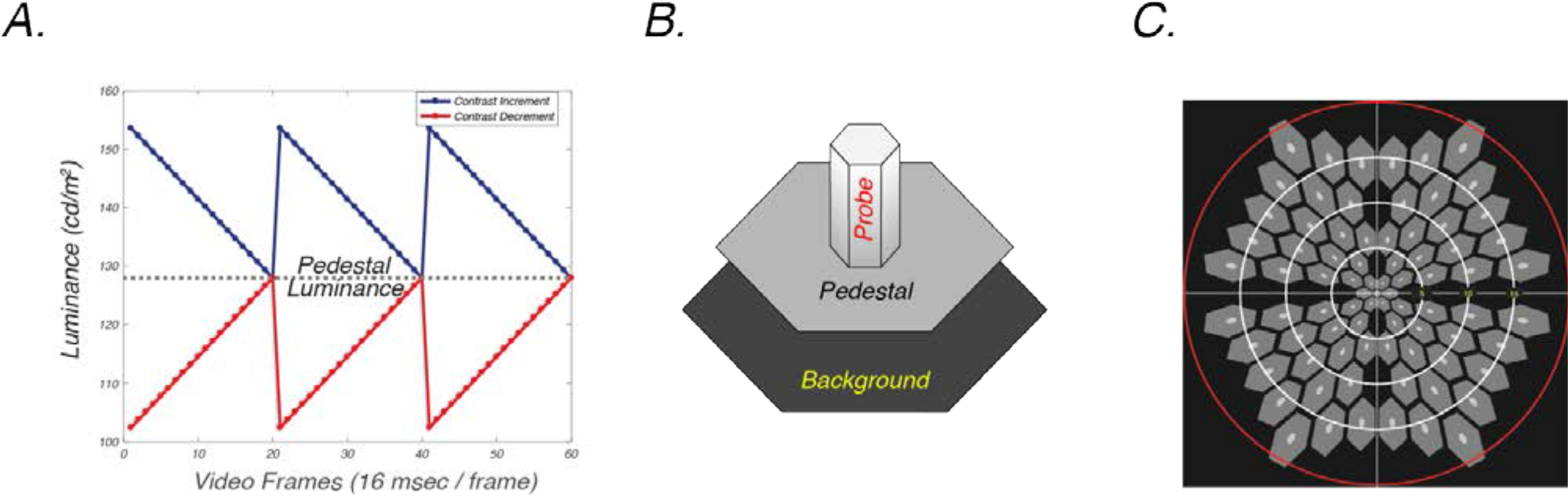
**A.** Probe waveforms. Incremental (blue) and decremental (red) sawtooth waveforms designed to favor ON vs OFF pathway responses, respectively. A stimulus frequency of 3 Hz is illustrated. **B.** Probe on pedestal display element. The sawtooth-modulated probes (small white hexagon) were presented on a mid-gray pedestal (medium size hexagon. An incremental pedestal is illustrated. The probe was 20% the size of the pedestal. The pedestal was surrounded by a black background region (largest hexagon). Weber contrast was 20% for both increments and decrements. **C.** Scaled stimulus array. The visual field was tiled with a set of probe/pedestal elements. The size of the elements was scaled over eccentricity according to the cortical magnification factor to optimize responses from the periphery. Typical field size was ~20 deg in radius (rings indicate 5 deg eccentricity radii from central fixation).

### General Procedure

The stimuli were presented in trials that lasted 12 *sec* for Experiment 1, 11 sec and 13.2 *sec* for the Experiments 2 and 3, respectively. The first and last seconds (Experiment 1) and the first and last 1.1 seconds for Experiments 2 and 3 were excluded from the data analysis. Participants were instructed to withhold blinking and fixate on the center element in Experiments 1 and 3. In Experiment 2, the participants performed a concurrent letter discrimination task presented within the central hexagon designed to control fixation and attention. The stimulus conditions were presented in random order with 3000 *msec* inter-trial intervals. Viewing was monocular in each experiment.

### Experiment 1 visual stimuli and participants

This experiment compared incremental and decremental responses for five stimulus configurations (see Fig. 3, top panels). One stimulus configuration comprised a rectangular array of 100 elements with the probe being 10.5 by 10.5 *arc min*, the pedestal 136 by 136 *arc min*. The entire array subtended 23.8 by 23.8 *deg* when viewed at 70 *cm*. Four additional configurations comprised hexagonal arrays of probe-on-pedestal elements. These arrays also covered 23.8 by 23.8 *deg* and were viewed at 70 cm. The base element in the center of the visual field varied between 28 and 80 *arc min* for the pedestal and between 5.6 to 16 *arc min* for the corresponding probes. The size of the array elements was scaled based on prior estimates of the cortical magnification factor (Baseler et al., 1994). The probe, pedestal and background luminance values were the same as for the rectangular array described above. This experiment was conducted with 19 participants (10 female, mean age 25). The experiment comprised 10 conditions (5 spatial layouts and 2 contrast signs). Nine trials lasting 12 *sec* were collected and the order of presentation was randomized.

### Experiment 2 visual stimuli and participants

This experiment was designed to collect data from a larger sample of participants with a display that eliminated elements that straddled the horizontal and vertical meridian (see Fig. 1C). This experiment utilized an 8 *arcmin* probe/40 *arcmin* pedestal display and was conducted with 30 participants (11 female, mean age 20). Nine trials lasting 13.2 sec were presented for both increments and decrements (8 conditions).

### Experiment 3 visual stimuli and participants

This experiment was designed to measure the relative amplitudes of the response in the upper and lower visual fields and to test for field cancellation effects that may have been present in the full-field recordings. This experiment utilized an 8 *arcmin* probe/40 *arcmin* pedestal display and was conducted with 17 participants (8 female, mean age 20). Eighteen trials lasting 11 sec were presented for both increments and decrements, each presented in both the upper and lower visual field (4 conditions).

### SSVEP recording

The EEG was recorded over 128 channels using Hydrocell SensorNets and NetStation 5.2 software (Electrical Geodesics, Eugene, OR). Prior to recording, individual electrodes were adjusted so that the impedance values were lower than 60 kΩ.

### Artifact rejection and EEG filtering

The raw EEG was amplified (gain = 1000 at 24-bit resolution) and digitally filtered with a zero-phase 0.3 Hz to 50 Hz bandpass filter. The data was then processed using in-lab software written in C++. The artifact rejection procedure first detected and then substituted consistently noisy individual channels. The noisy channels were substituted them with the average signals of the six nearest electrodes surrounding the noisy electrode. After this, the EEG was re-referenced to the common average of all electrodes. Secondly, in an effort to reject data recorded during coordinated muscle movements or blinks, 1-second-long epochs were excluded for all electrodes if signals of more than 5% (7 out of 128) of the electrodes exceeded a set threshold amplitude (60-520 µV, median: 100 µV) sometime during the epoch. Finally, 1 sec epochs from individual electrodes were excluded if more than 10% of the epoch samples exceeded +/− 30 µV.

### Reliable Components Analysis

Reliable Components analysis (RCA; (Dmochowski et al., 2015) was used to reduce the dimensionality of the 128 channel recordings to a small number of components. Each RCA component comprises a scalp topography and a response time-course or spectrum. RCA components were derived through an eigenvalue decomposition that maximized the trial-by-trial covariance matrix. This criterion reflects the fundamental assumption that the stimulus-driven evoked response is highly similar over repeated presentations of the same stimulus. RCA also results in an improvement signal-to-noise ratio and provides a data driven method for selecting the recording channels whose data are to be analyzed. RCA was computed over conventional time-averages over the cycle length of the stimulus which was 333 msec in Experiment 1, and 366 msec for Experiments 2 and 3, or over the complex values of the first 4 harmonics of the stimulus frequency (Dmochowski et al., 2015)

## Results

### Experiment 1: Scaling the stimulus array to maximize response amplitude

The VEP stimuli were comprised of multiple small elements because our ultimate goal is to probe ON *vs* OFF pathway responses in different parts of the visual field. Prior work with the isolated check VEP has used rectangular arrays of small probes on a large, uniform intensity background (Zemon et al., 2008; Chen and Zhao, 2017; Xu et al., 2017(Greenstein et al., 1998). Other work on the multi-focal VEP, by contrast, has scaled the elements of multi-element displays in order to equate the cortical area being stimulated over retinal eccentricity (Baseler et al., 1994). This has the benefit of roughly equating SNR for elements presented at different locations in the visual field. To determine whether scaling the VEP test/pedestal stimulus for retinal eccentricity results in a larger VEP than an unscaled stimulus, we compared responses generated by several cortically scaled arrays to those from an unscaled array similar to one used in previous experiments on incremental and decremental SSVEP responses (Zemon et al., 2008; Chen and Zhao, 2017; Xu et al., 2017; (Greenstein et al., 1998).

The logic of the scaling is most easily understood by considering the central element of the multi-element array. This element is at the point of fixation (the fovea) where cortical magnification is highest. We sought to determine what the optimal spatial scale for the entire array would be to elicit the largest VEP. This scale would presumably be the one that drives the largest number of cells at each eccentricity. Because of cortical magnification, the probe and pedestal in the fovea should be much smaller than the probes and pedestals in the periphery. We reasoned that by fixing the cortical magnification factor and varying the size of central element, we could determine which set of scaled elements resulted in the largest evoked response. After some preliminary testing, we settled on a range of element sizes that spanned a factor of 4 in half-octave steps. The results of Experiment 1 are shown as time averages in Figure 3 for both decrements (red/orange traces) and increments (blue/cyan traces). The largest responses measured were for decrements at a central pedestal size of 40 arcmin (8 arcmin probe size). Responses to increments were smaller and did not differ between 28 and 40 arcmin scaled arrays, but were lower at 56 and 80 arcmin. Responses to decrements were thus more spatially tuned, being smaller for both the 28 and 56 arcmin stimuli. For each of the scaled arrays, the latency of the negative peak is slightly shorter for decrements than for increments.

**Fig. 2.**
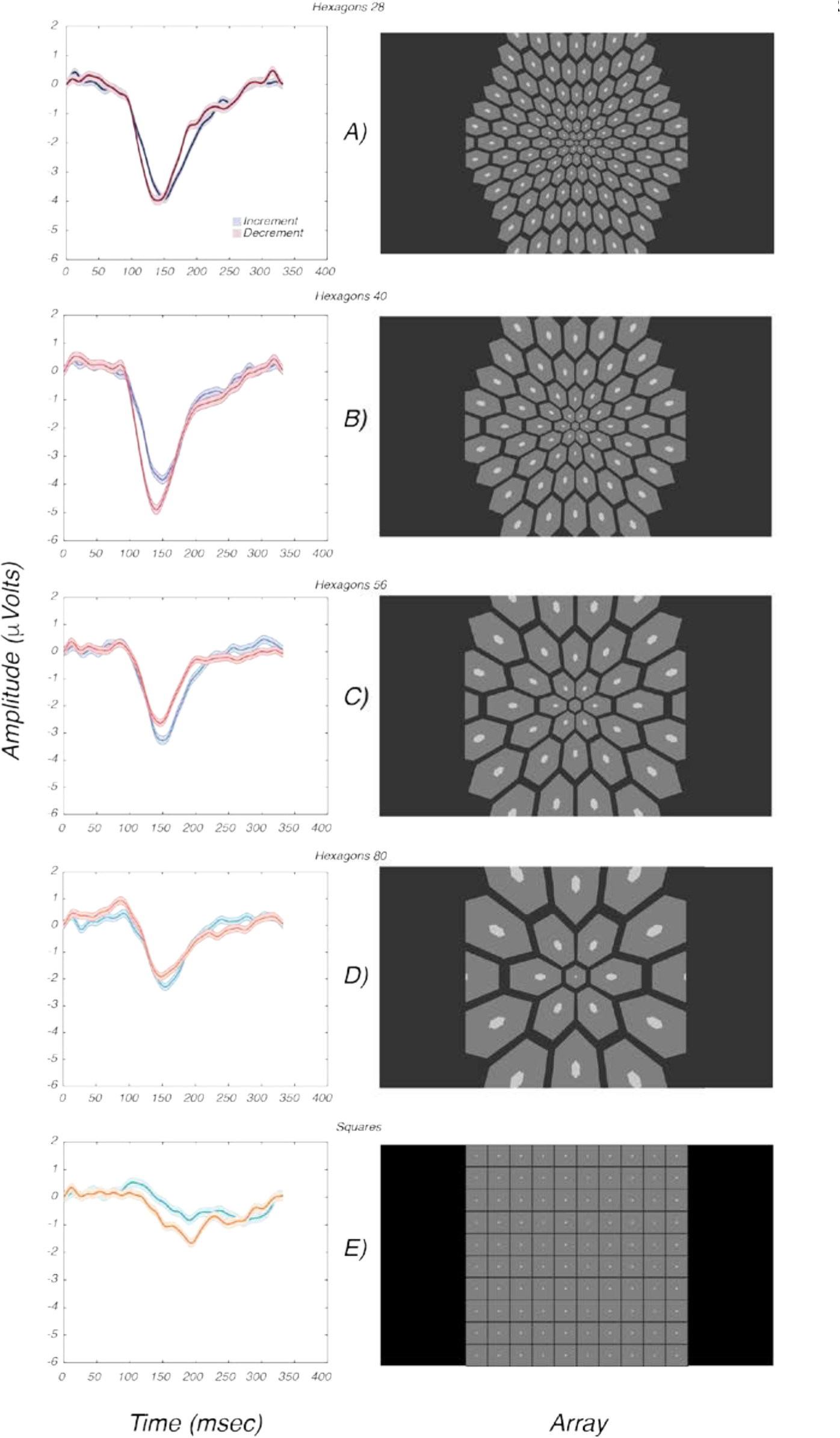
Effect of varying element scaling and format. A)-E) Increment (blue) and decrement (red) responses (left panels) for different array scalings and formats (right panels).

Figure 3E shows the data from the rectangular array. Both increment and decrement responses are smaller than any of the responses measured with scaled arrays. This is despite there being more probes in the rectangular array (100) than in 3 of 4 of the scaled arrays (125, 59, 35 and 19 respectively). The response to decrements is larger for the rectangular array, as it was for the hexagonal display scaling that resulted in the largest overall response (40 arc min). Given that the 40 arcmin scaled array evoked the largest response and that the response differed for increments and decrements, this display scaling was used for the remaining experiments.

**Fig. 3.**
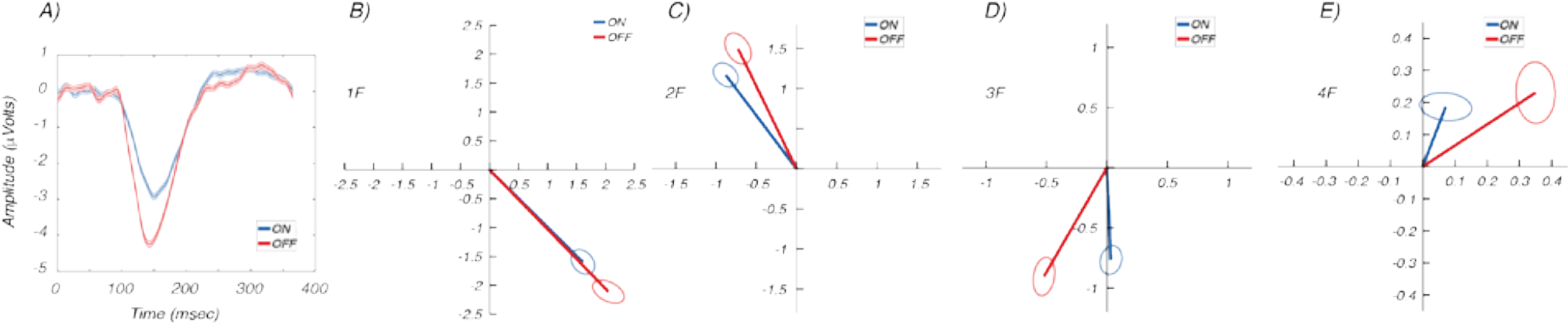
A. Evoked response in the time-domain for full-field increments (blue) and decrements (red). Error bands plot +/− 1 s.e.m. Response is larger and latency shorter for decrements (n=30 eyes). B-E. Evoked response in the frequency domain at 1F, 2F, 3F and 4F, respectively. Responses to increments are in blue, decrements in red. Error ellipses represent 1 s.e.m contours.

### Experiment 2. Full-field responses with modified array

In this experiment, we used the 40 arcmin scaled array to collect monocular data for incremental and decremental stimuli in a group of 30 participants. Cycle-averaged waveforms for increments are shown in Figure 4A as the blue trace and for decrements as the red trace in Fig. 3A. The time domain waveforms in this group of participants were similar to those shown in Figure 3B for the same element size of 40 arcmin, with responses to decrements being ~1.4 times larger than those for increments. There was also an ~ 5 msec difference in the time-to-peak of the dominant negative-going response component, with decrement responses being faster than increment responses.

**Fig. 4.**
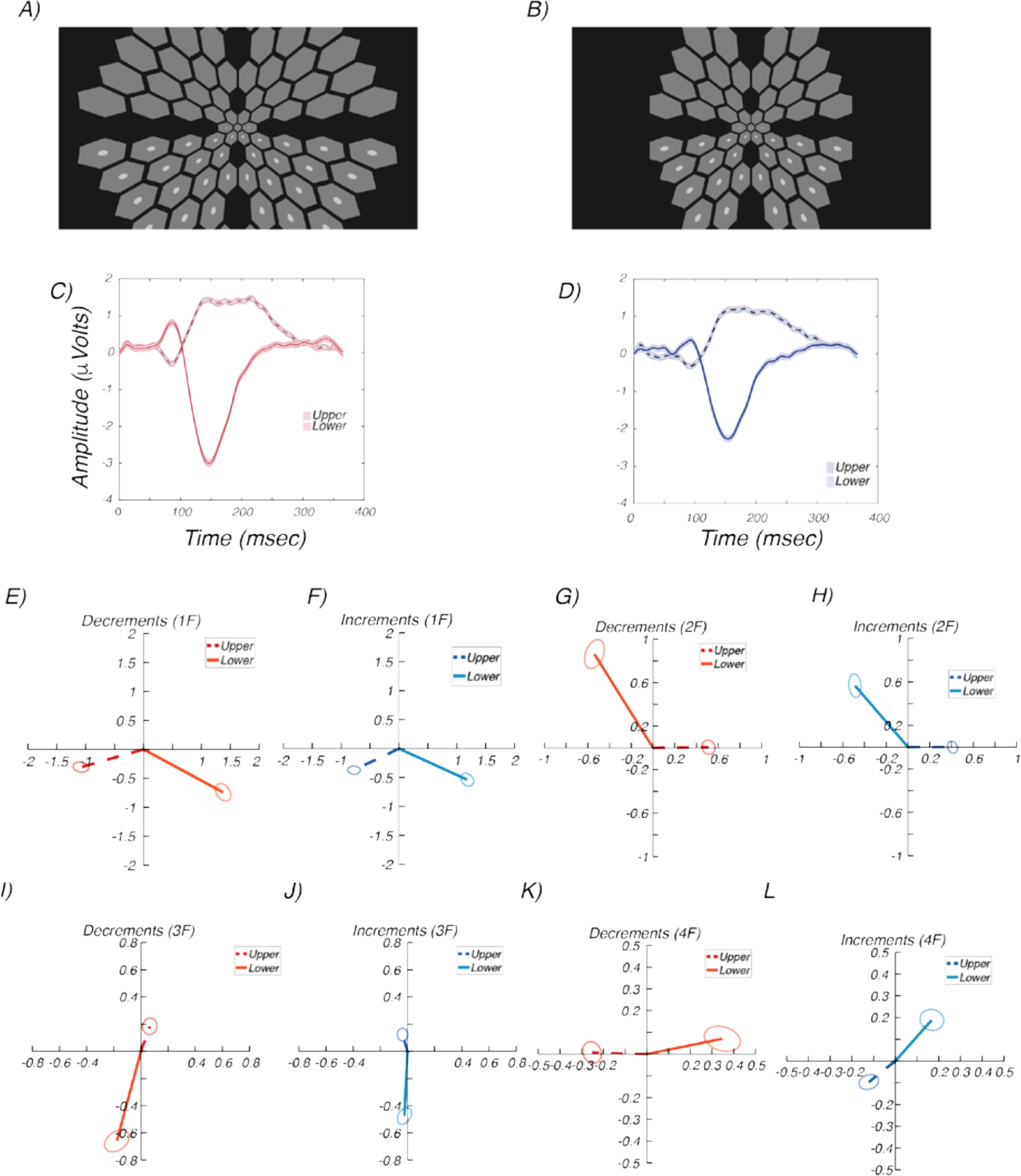
Upper vs lower visual field responses. A) Example large-field display (lower field/increments) B) Example smaller-field display (lower field/increments). Both upper and lower and incremental and decremental displays were used. C) Response waveform, upper vs lower field increments D) Response waveform, upper vs lower field increments E-L Upper field (dashed) vs lower field (solid) responses for increments (warm colors) and decrements (cool colors) for 1F, 2F, 3F and 4F response components. Responses to lower field are larger than for upper field. Responses are approximately polarity inverted between hemifields. Decrement responses are larger and faster than increment responses. See text for details.

Figure 3 B-E shows the same data in the frequency domain for the first four harmonics of the 2.73 Hz stimulus frequency. The response to decrements is larger than that for increments for all 4 harmonics and the phase of the response shows less delay at the 2^nd^, 3^rd^ and 4^th^ harmonics for decrements, consistent with the earlier peak latency for decrements seen in Figure 3A.

### Experiment 3: Upper vs lower visual field response properties

The three-dimensional geometry of early retinotopic visual areas V1, V2 and V3 is such that source orientations in the upper and lower visual fields lead to polarity reversals at the scalp and thus to potential field cancellations when both upper and lower fields are stimulated at the same time (Ales et al., 2010). Moreover, previous research has found that lower field responses are typically larger than upper field responses (Skrandies, 1987; Hagler, 2014). To determine the extent to which our responses were affected by field cancellation and to determine the relative magnitude of upper and lower field responses, we repeated the measurements with separate upper and lower field stimulation runs.

Upper and lower field responses for two stimulus arrays that differed in visual field extent (Fig. 4 A and B) were combined and are shown as cycle averages in Fig. 4C and D and as frequency components in Fig. 4E-4L. In the time-domain, there is an initial lower visual field positivity near 85 msec for decrements and a corresponding upper-field negativity at the same latency (compare Fig. 4C solid and dashed, curves, respectively. These matching negative/positive peaks are also present for increments, but the peak latency is later at approximately 95 msec (Fig. 4D solid *vs* dashed curves). The polarity inversion of the 85/95 msec peak is consistent with sources in early visual cortex (Ales et al., 2010). Within the lower visual field, responses to decrements are larger than those for increments by a factor of 2.5 for the 85/95 msec peak (compare solid curves in Fig. 4C to those of Fig. 4D). The corresponding responses in the upper field are also larger for decrements, but by a smaller fraction at this latency. The response amplitudes at 85 msec, the time of the initial peak for decrements, differ by factor ~4.5 between upper and lower field responses with lower field responses being larger (see Fig. 4C). For the ~95 msec peak for increments, lower field responses are larger by a factor of ~1.8 (see Fig. 4D).

These initial deflections are followed by larger peaks, consisting of a ~150 msec negativity for both decrements and increments in the lower visual field. In the upper visual field, the potential comprises a broader positivity of different. The mid-latency lower field responses are also larger than the corresponding upper field responses for both decrements and increments (a factor of ~ 2 for both decrements and increments). The decremental response peaks occur ~ 10 msec earlier than the increment responses at both early (85/90 msec) and mid latencies (~150 msec) and these latency differences are of the same magnitude in the upper and lower visual fields. As noted, decrement responses have larger peak amplitudes at both ~90 and ~150 msec in the lower visual field. In the upper visual field, the ~90 msec negative peak is larger but the later positivity is the same amplitude at mid latencies.

Responses for 1F, 2F, 3F and 4F response components are shown in Fig 4E-5L. Here the large amplitude differences between lower and upper visual field are again apparent in the length of the response vectors. The shorter latency for decrements than for increments is also apparent in the frequency-domain results where it manifests as phase leads.

To quantify the magnitude of the timing difference between increments and decrements, we derived apparent latencies in the frequency-domain by plotting response phase as a function of response frequency (Lopes da Silva et al., 1970; Yeatman and Norcia, 2016). Delay was estimated as d = 1/360 X δϕ/δf, where δϕ is the change in phase over the frequency range in degrees and δf is the change in frequency in Hz.

The measured response phase is a linear function of component frequency, consistent with a constant delay. The estimated delays are 135.6 +/−0 .15 and 146.0 +/−0.08 msec for decrements and increments, respectively in the lower visual field (Fig. 5, left) and 111.6 +/−0.36 and 124.66 +/− 0.21 for the corresponding stimulus polarities for the upper visual field (Fig. 5, right). The latency advantage for decrements is thus 8.4 msec in the upper visual field and 15.06 msec in the lower visual field.

**Fig. 5.**
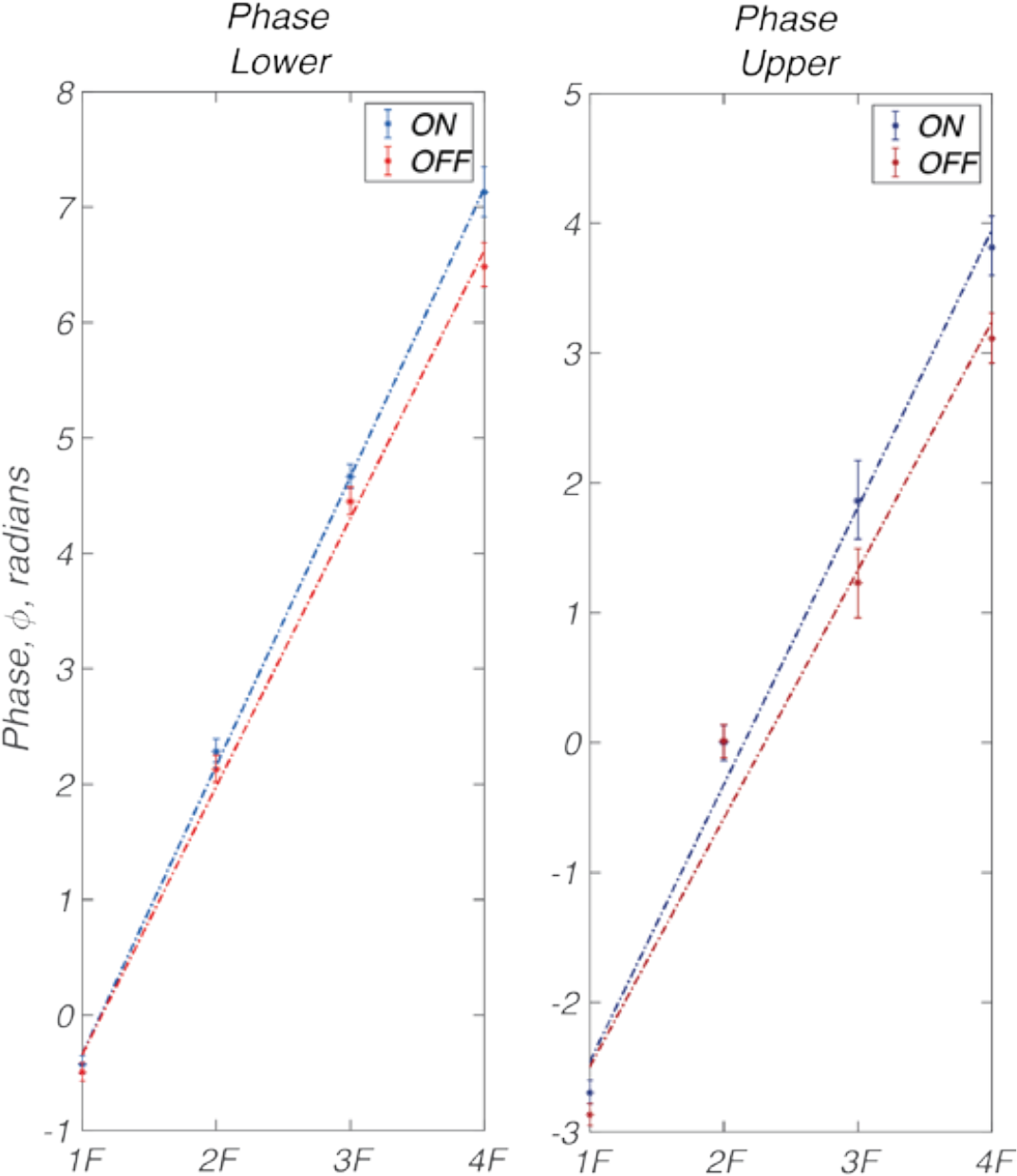
Response phase as a function of frequency (harmonicnumber). Left panel plots increment (OFF, blue) and decrement (ON, red) response phases for the lower visual field. Right panel uses the same conventions to plot data from the upper visual field. Linear regression lines are fit to the data to estimate delays (see text for details). The derived latency advantage for decrements is thus 8.4 msec in the upper visual field and 15.06 msec in the lower visual field.

## Discussion

Here we find that VEPs to equal-value sawtooth increments versus decrements recorded at the scalp differ in several respects. The spatial tuning for decrements is narrower than for increments and responses to decrements are typically larger and faster than responses to increments. In addition, we found lower field responses to be substantially larger than upper field responses.

### Comparison with previous VEP studies of increment and decrement responses

Several previous studies have compared VEPs to contrast increments and decrements. Zemon et al., (1988) introduced the use of isolated checks on a mid-gray background to allow for separate testing of contrast increments and decrements. The checks were sinusoidally modulated in luminance above and below the mid-level luminance of a static background to bias responses to the ON *vs* OFF pathways, respectively. In the single-subject records shown in the paper, responses to decrements were consistently larger than those for increments, a pattern we find to typically hold, as well. They also reported that the spatial tuning for incremental *vs* decremental stimuli depended on the size of the checks, being consistently larger for smaller, decremental checks than for corresponding incremental checks of the same size. Responses to our rectangular-array stimulus showed a strong amplitude bias in favor of decrements under probe-size conditions that produced a consistently large bias in favor of decrements in Zemon et al’s measurements. We also found that the relative amplitude of incremental *vs* decremental stimuli depends on the spatial parameters of the display, but we found that spatial tuning was narrower for decrements, rather than for increments, as was the case in Zemon et al., 1988). A later study working within the same paradigm used sinusoidally modulated isolated checks presented at 6 and 15 Hz (Greenstein et al., 1998). Although not specifically analyzed, it is apparent from their Figure 5 that the amplitudes for increments and decrements were very similar in their normal vision control group, while we find larger responses for decrements.

Other considerably smaller studies that have compared incremental decremental response have found mixed results. A study using transient onset/offset of contrast increments and decrements found larger increment responses in a group of 5 participants (Mutlukan et al., 1992). Another study used 2 Hz saw-tooth waveforms similar to ours and found no amplitude differences between incremental *vs* decremental stimuli in two participants (Roveri et al., 1997). Their use of sawtooth waveforms, as ours was motivated by the finding the fast increment/slow decrement waveforms preferentially activated ON retinal ganglion cell in the macaque while fast decrement/slow increment waveforms activation OFF cells (Kremers et al., 1993). Some of the difference between our study and these previous ones could have been due to sampling biases in small studies. We find larger amplitudes for decrements over three studies involving 68 participants. Nonetheless, considering our data in Fig. 3 and the data from previous studies reviewed above, it appears that the relative amplitude of incremental *vs* decremental responses may depend on the conditions of measurement and may not be a fixed property of all stimuli.

In terms of the dynamics of ON and OFF pathways, Zemon et al., (1988) and Greenstein et al (1998) both measured the phase of the Steady-State VEP for increments and decrements (typical response frequency of 6 Hz) and found no measurable differences for the two contrast polarities. Similarly, no latency differences were reported in the other studies that have compared incremental and decremental responses (Mutlukan et al., 1992; Roveri et al., 1997). In contrast to these previous studies, we find small, but measurable timing differences between decremental and incremental responses. The differences are most readily apparent in the hemifield data, where we could measure both early latency and mid latency components in the time domain and corresponding phase shifts across lower and higher harmonic responses. These differences were small, making them difficult to measure. Moreover, the use of full field stimulation, as in previous studies may complicate the measurement. Responses from early visual areas may be obscured with full-field recordings (compare response amplitude at ~90 msec in Fig. 4 to that of Fig. 5, for example).

### Upper vs lower field responses

We found that lower field responses were a factor of ~ 2 larger than upper field responses, something that has been consistently observed in previous studies (Skrandies, 1987; Hagler, 2014). Also consistent with previous work is our finding that response components invert polarity between upper and lower visual fields (Jeffreys and Axford, 1972a, b; Di Russo and Spinelli, 2002; Di Russo et al., 2005). This polarity inversion is only approximate however. In the time domain, the shape of the upper and lower field waveform differs substantially (see Fig. 5). In the frequency domain, the upper and lower field responses differ in phase by between ~90 deg (1F increments) to ~120 deg at 2F rather than the 180 degrees that would be expected from pure geometry-based polarity inversion. By contrast, responses at 3F and 4F are phase shifted by ~180 deg. This suggests that 3F and 4Fresponses are dominated by early visual cortex (V1,V2 and V3) where the aggregate tissue orientation of the upper and lower visual field representations can cause scalp polarity inversions (Ales et al., 2010). The 1F and 2F responses may contain contributions from higher visual areas, such as V3A and V4 that have complete hemifield representations, rather than the split dorsal (lower visual field) and ventral (upper visual field) representations of V1, V2 and V3.

### Comparison with single-unit data

Our finding of generally larger response amplitudes for decremental sawtooth stimuli is consistent with the OFF cell bias in cortex that has been found in single-unit studies from a variety of species (Jin et al., 2008; Yeh et al., 2009; Xing et al., 2010; Kremkow et al., 2014; Liu and Yao, 2014; Zurawel et al., 2014).

Several in vivo studies have reported that OFF cells have shorter latency than ON cells in the LGN (Jin et al., 2011b; Li et al., 2017) and in visual cortex (Komban et al., 2014; Jiang et al., 2015; Rekauzke et al., 2016), a pattern we have also observed. These latency differences are small, being 8-15 msec (Fig. 5) in our data and between 3-6 msec in cat LGN (Jin et al., 2011b; Li et al., 2017),3 msec in cat visual cortex (Komban et al., 2014) and 5 msec in macaque V1 measured with voltage-sensitive dye imaging (Rekauzke et al., 2016). Note that one earlier in vitro study (Chichilnisky and Kalmar, 2002) however, found the opposite. The failure of previous VEP studies to detect small magnitude latency differences may be been due to their limited statistical power or the inherent sensitivity of the different measurement approaches.

OFF biases seen in the single unit data have been argued to underlie the lower detection thresholds and faster reaction times that have been reported for contrast decrements (Komban et al., 2014). Our data lend additional support to these suggestions. The amplitude biases we observe may arise in cortex, given that individual ON and OFF cells in the retina and LGN have not been reported to differ in their sensitivity (Kremers et al., 1993). The latency biases we observe may arise sub-cortically and be passed on to cortex where they appear as a constant temporal offset at all early and mid-latencies.

### Conclusion

In summary, we find that the VEP can be used to discriminate ON *vs* OFF visual pathway signaling in human, corroborating several previously observed response biases measured in single cells in animal models. Future directions include exploring amplitude and timeing relationships of the ON and OFF pathways as a function of visual field location, subject age, and presence of visual system disease such as has been reported in animal models of glaucoma (Della Santina et al., 2013; El-Danaf and Huberman, 2015; Ou et al., 2016; Puyang et al., 2017; Daniel et al., 2018).

## References

Ales JM, Yates JL, Norcia AM (2010) V1 is not uniquely identified by polarity reversals of responses to upper and lower visual field stimuli. Neuroimage 52:1401–1409.

Baseler HA, Sutter EE, Klein SA, Carney T (1994) The topography of visual evoked response properties across the visual field. Electroencephalogr Clin Neurophysiol 90:65–81.

Chichilnisky EJ, Kalmar RS (2002) Functional asymmetries in ON and OFF ganglion cells of primate retina. J Neurosci 22:2737–2747.

Dacey DM, Petersen MR (1992) Dendritic field size and morphology of midget and parasol ganglion cells of the human retina. Proc Natl Acad Sci U S A 89:9666–9670.

Del Viva MM, Gori M, Burr DC (2006) Powerful motion illusion caused by temporal asymmetries in ON and OFF visual pathways. J Neurophysiol 95:3928–3932.

Devries SH, Baylor DA (1997) Mosaic arrangement of ganglion cell receptive fields in rabbit retina. J Neurophysiol 78:2048–2060.

Di Russo F, Spinelli D (2002) Effects of sustained, voluntary attention on amplitude and latency of steady-state visual evoked potential: a costs and benefits analysis. Clin Neurophysiol 113:1771–1777.

Di Russo F, Pitzalis S, Spitoni G, Aprile T, Patria F, Spinelli D, Hillyard SA (2005) Identification of the neural sources of the pattern-reversal VEP. Neuroimage 24:874–886.

Dmochowski JP, Greaves AS, Norcia AM (2015) Maximally reliable spatial filtering of steady state visual evoked potentials. Neuroimage 109:63–72.

Dowling JE, Werblin FS (1971) Synaptic organization of the vertebrate retina. Vision Res Suppl 3:1–15.

Famiglietti EV, Jr., Kolb H (1976) Structural basis for ON-and OFF-center responses in retinal ganglion cells. Science 194:193–195.

Geisler WS (2008) Visual perception and the statistical properties of natural scenes. Annu Rev Psychol 59:167–192.

Greenstein VC, Seliger S, Zemon V, Ritch R (1998) Visual evoked potential assessment of the effects of glaucoma on visual subsystems. Vision Res 38:1901–1911.

Hagler DJ, Jr. (2014) Visual field asymmetries in visual evoked responses. J Vis 14:13.

Jeffreys DA, Axford JG (1972a) Source locations of pattern-specific components of human visual evoked potentials. II. Component of extrastriate cortical origin. Exp Brain Res 16:22–40.

Jeffreys DA, Axford JG (1972b) Source locations of pattern-specific components of human visual evoked potentials. I. Component of striate cortical origin. Exp Brain Res 16:1–21.

Jiang Y, Purushothaman G, Casagrande VA (2015) The functional asymmetry of ON and OFF channels in the perception of contrast. J Neurophysiol 114:2816–2829.

Jin J, Wang Y, Swadlow HA, Alonso JM (2011a) Population receptive fields of ON and OFF thalamic inputs to an orientation column in visual cortex. Nat Neurosci 14:232–238.

Jin J, Wang Y, Lashgari R, Swadlow HA, Alonso JM (2011b) Faster thalamocortical processing for dark than light visual targets. J Neurosci 31:17471–17479.

Jin JZ, Weng C, Yeh CI, Gordon JA, Ruthazer ES, Stryker MP, Swadlow HA, Alonso JM (2008) On and off domains of geniculate afferents in cat primary visual cortex. Nat Neurosci 11:88–94.

Komban SJ, Kremkow J, Jin J, Wang Y, Lashgari R, Li X, Zaidi Q, Alonso JM (2014) Neuronal and perceptual differences in the temporal processing of darks and lights. Neuron 82:224–234.

Kremers J, Lee BB, Pokorny J, Smith VC (1993) Responses of macaque ganglion cells and human observers to compound periodic waveforms. Vision Res 33:1997–2011.

Kremkow J, Jin J, Wang Y, Alonso JM (2016) Principles underlying sensory map topography in primary visual cortex. Nature 533:52–57.

Kremkow J, Jin J, Komban SJ, Wang Y, Lashgari R, Li X, Jansen M, Zaidi Q, Alonso JM (2014) Neuronal nonlinearity explains greater visual spatial resolution for darks than lights. Proc Natl Acad Sci U S A 111:3170–3175.

Lee KS, Huang X, Fitzpatrick D (2016) Topology of ON and OFF inputs in visual cortex enables an invariant columnar architecture. Nature 533:90–94.

LeVay S, McConnell SK (1982) ON and OFF layers in the lateral geniculate nucleus of the mink. Nature 300:350–351.

Li H, Liu X, Andolina IM, Li X, Lu Y, Spillmann L, Wang W (2017) Asymmetries of Dark and Bright Negative Afterimages Are Paralleled by Subcortical ON and OFF Poststimulus Responses. J Neurosci 37:1984–1996.

Liu K, Yao H (2014) Contrast-dependent OFF-dominance in cat primary visual cortex facilitates discrimination of stimuli with natural contrast statistics. Eur J Neurosci 39:2060–2070.

Lopes da Silva FH, van Rotterdam A, Storm van Leeuwen W, Tielen AM (1970) Dynamic characteristics of visual evoked potentials in the dog. II. Beta frequency selectivity in evoked potentials and background activity. Electroencephalogr Clin Neurophysiol 29:260–268.

McConnell SK, LeVay S (1984) Segregation of on- and off-center afferents in mink visual cortex. Proc Natl Acad Sci U S A 81:1590–1593.

Mutlukan E, Bradnam M, Keating D, Damato BE (1992) Visual evoked cortical potentials from transient dark and bright stimuli. Selective ‘on’ and ‘off-pathway’ testing? Doc Ophthalmol 80:171–181.

Nichols Z, Nirenberg S, Victor J (2013) Interacting linear and nonlinear characteristics produce population coding asymmetries between ON and OFF cells in the retina. J Neurosci 33:14958–14973.

Ratliff CP, Borghuis BG, Kao YH, Sterling P, Balasubramanian V (2010) Retina is structured to process an excess of darkness in natural scenes. Proc Natl Acad Sci U S A 107:17368–17373.

Rekauzke S, Nortmann N, Staadt R, Hock HS, Schoner G, Jancke D (2016) Temporal Asymmetry in Dark-Bright Processing Initiates Propagating Activity across Primary Visual Cortex. J Neurosci 36:1902–1913.

Roveri L, Demarco PJ, Jr., Celesia GG (1997) An electrophysiological metric of activity within the ON- and OFF-pathways in humans. Vision Res 37:669–674.

Skrandies W (1987) The upper and lower visual field of man: electrophysiological and functional differences. Prog Sensory Physiol 8:1–93.

Smith GB, Whitney DE, Fitzpatrick D (2015) Modular Representation of Luminance Polarity in the Superficial Layers of Primary Visual Cortex. Neuron 88:805–818.

Stryker MP, Zahs KR (1983) On and off sublaminae in the lateral geniculate nucleus of the ferret. J Neurosci 3:1943–1951.

Wang Y, Jin J, Kremkow J, Lashgari R, Komban SJ, Alonso JM (2015) Columnar organization of spatial phase in visual cortex. Nat Neurosci 18:97–103.

Wassle H (2004) Parallel processing in the mammalian retina. Nat Rev Neurosci 5:747–757.

Westheimer G (1965) Spatial interaction in the human retina during scotopic vision. J Physiol 181:881–894.

Westheimer G (1967) Spatial interaction in human cone vision. J Physiol 190:139–154.

Xing D, Yeh CI, Shapley RM (2010) Generation of black-dominant responses in V1 cortex. J Neurosci 30:13504–13512.

Yeatman JD, Norcia AM (2016) Temporal Tuning of Word- and Face-selective Cortex. J Cogn Neurosci 28:1820–1827.

Yeh CI, Xing D, Shapley RM (2009) “Black” responses dominate macaque primary visual cortex v1. J Neurosci 29:11753–11760.

Zaghloul KA, Boahen K, Demb JB (2003) Different circuits for ON and OFF retinal ganglion cells cause different contrast sensitivities. J Neurosci 23:2645–2654.

Zahs KR, Stryker MP (1988) Segregation of ON and OFF afferents to ferret visual cortex. J Neurophysiol 59:1410–1429.

Zemon V, Gordon J, Welch J (1988) Asymmetries in ON and OFF visual pathways of humans revealed using contrast-evoked cortical potentials. Vis Neurosci 1:145–150.

Zemon V, Eisner W, Gordon J, Grose-Fifer J, Tenedios F, Shoup H (1995) Contrast-dependent responses in the human visual system: childhood through adulthood. Int J Neurosci 80:181–201.

Zurawel G, Ayzenshtat I, Zweig S, Shapley R, Slovin H (2014) A contrast and surface code explains complex responses to black and white stimuli in V1. J Neurosci 34:14388–14402.

